# Microbial Modulation of Host Plant Proteasome Activity Improves Heat Stress Tolerance

**DOI:** 10.1101/2025.05.27.656411

**Authors:** Jun Hyung Lee, Alyssa A. Carrell, Bryan Piatkowski, Dana L. Carper, Lee Gunter, Sara Jawdy, Kelsey Carter, Mitchel J. Doktycz, Paul E. Abraham, Jeremy Schmutz, John Lovell, Adam Healey, Dale A. Pelletier, David J. Weston

**Affiliations:** Biosciences Division, Oak Ridge National Laboratory, 1 Bethel Valley Rd., Oak Ridge, TN 37831, USA; Department of Environmental Biology, State University of New York College of Environmental Science and Forestry, 1 Forestry Dr., Syracuse, NY 13210; Environmental Sciences Division, Oak Ridge National Laboratory, 1 Bethel Valley Rd., Oak Ridge, TN 37831, USA; HudsonAlpha Institute for Biotechnology, 601 Genome Way, Huntsville, AL 35806, USA; Department of Energy Joint Genome Institute, Lawrence Berkeley National Laboratory, 1 Cyclotron Rd., CA 94720, USA

**Keywords:** ECM29, endophyte, sphagnum, thermotolerance, proteasome

## Abstract

Global climate change marked by rising temperatures and increasingly frequent, severe heat waves, threatens plant health, productivity, and ecosystem function. Beneficial plant-microbe associations are increasingly recognized for their role in mitigating abiotic stress, yet the molecular mechanisms underlying microbially-conferred thermotolerance remain poorly understood. Here, we developed a population-based assay using *Sphagnum* peat mosses, a key species in peatland ecosystems that sequester 33–50% of the world’s soil carbon as recalcitrant peat, to perform microbiome transfers and assess microbial contributions to heat resilience. Strain-based assays in *Sphagnum* and *Arabidopsis* identified *Variovorax* as a key thermotolerance-enhancing microbe and identified ECM29, a proteasome-associated protein, as critical for this interaction. In *Arabidopsis, Variovorax* sp. CF313 association led to ECM29 preferentially binding the RPT1 subunit of the 19S regulatory particle, inhibiting 20S proteasome gate opening and slightly reducing proteasomal activity. This results in the accumulation of ubiquitinated proteins and induces *HSP70* transcription, priming the host for heat shock stress even in the absence of prior heat exposure. These findings establish ECM29 as a key regulator of proteasome activity in response to microbial interactions, highlighting a previously unknown mechanism by which microbes enhance plant thermotolerance. More broadly, this work highlights the importance of leveraging natural systems and ecologically relevant models to identify strategies for enhancing plant and ecosystem resilience to a changing climate.

**Significance Statement:** Global climate change poses severe threats to plant health and productivity, with cascading impacts on ecosystems. Beneficial plant-microbe interactions offer promising strategies to mitigate abiotic stress, yet the molecular mechanisms often remain unclear. This study uncovers a novel thermotolerance mechanism mediated by *Variovorax* modulating the plant host proteasome. Using a QTL derived from *Sphagnum* peat mosses, and tested within a genetically tractable *Arabidopsis* model, we identified ECM29, a highly conserved protein across diverse plant species, as a key regulator of proteasome activity. ECM29 primes plants for heat stress by inducing HSP70 transcription through proteasome modulation. Additionally, we demonstrated that ECM29 is required for the host to receive microbially conferred thermotolerance, but is not necessary for innate priming to heat stress. These findings highlight the potential of leveraging natural systems and ecologically relevant species to develop innovative genetic models, advancing plant resilience and productivity in a warming world.

## Introduction

Microbial communities play a pivotal role in shaping plant responses to environmental stress, particularly in the face of escalating climate change pressures (1, 2). Research has shown that microbiomes endemic to stressful environments can enhance host plant tolerance to similar conditions (3, 4). These findings have prompted efforts to identify the host traits and underlying genetic factors that influence microbial community composition and function (5). These interactions are especially important in boreal and subarctic peatlands, which serve as critical global carbon (C) sinks by storing 33–50% of the world’s soil C as recalcitrant soil peat (6, 7). Within these ecosystems, *Sphagnum* peat mosses and their associated microbiomes are key contributors to productivity, N_2_-fixation, and broader nutrient cycling (8, 9). Recent studies have shown that peatland warming is influencing the community composition of the *Sphagnum* microbiome, N_2_-fixation and methane oxidation and abiotic stress tolerance (10–12). Given rising temperatures, extended heat waves, and more frequent extreme weather events (13, 14), plant associated microbial communities can undergo rapid shifts in composition and function, potentially shaping host thermal tolerance and becoming a critical factor in environmental stress acclimation and adaptation (15–17). Across multiple plant host species, microbial symbionts have been shown to enhance thermotolerance through various mechanisms, including the induction of heat-responsive genes, production of protective metabolites, and modulation of nutrient allocation and composition (18–20). However, despite these advances, we still lack a fundamental molecular genetic understanding of how beneficial microbes bypass host defense mechanisms, establish beneficial relationships, and enhance host plant stress tolerance.

Seminal studies have demonstrated that plant associated microbes endemic to harsh environments can confer analogous stress tolerance to host plants (3, 4, 21). Recently, this paradigm was extended to peatland ecosystems, where microbes associated with *Sphagnum* from a whole-ecosystem warming study were transferred to laboratory-grown, warming-naïve *Sphagnum* hosts and shown to enhance growth at elevated temperatures(22). This microbiome transfer approach provides a system to examine how plants with different genetic backgrounds interact with and benefit from microbial communities pre-conditioned to elevated temperatures. Notably, these studies observed an enrichment of *Variovorax* within the warming treated microbiome, suggesting a specialized role for this bacterium in modulating host thermal tolerance. Our recent study identified that several strains in the genus *Variovorax,* including *Variovorax* sp. CF313 (23), confer thermal benefit to *Arabidopsis* (24). Further research has highlighted additional plant benefits conferred by *Variovorax*, including its role in promoting growth and diverse abiotic stress tolerance. *Variovorax* maintains host root growth by suppressing the inhibitory effects of other bacteria present in the complex microbial community, particularly by manipulating auxin levels (25). Plants inoculated with *Variovorax* showed enhanced tolerance to drought and salt stresses, potentially through 1-aminocyclopropane-1-carboxylic acid deaminase activity which lowers stress ethylene levels (26–28). While these studies establish *Variovorax* as a beneficial microbe capable of enhancing plant stress tolerance, the host genetic factors are necessary to receive these microbial benefits and the interacting plant and microbial pathways mediating thermotolerance remain largely unknown.

To better understand the genetic mechanisms by which plants acquire microbial-conferred thermotolerance, we developed a population-based assay using *Sphagnum fallax*, performed microbiome transfers, and conducted strain-based assays, along with heterologous transformation in *Arabidopsis*, to investigate the genetic basis of microbially enhanced thermal benefits. Microbiomes associated with *Sphagnum* were collected from a whole-ecosystem peatland warming study (29) and transferred across a *Sphagnum* pedigree (196 sequenced members) and temperature treatments (4 levels), encompassing 2,365 unique plant-microbe-treatment combinations. This approach identified a seven-gene QTL region associated with host reception of microbially provided thermotolerance. To uncover the molecular mechanisms underlying this microbially provided thermotolerance, we focused on *Variovorax*, a bacterial genus enriched in the warming microbiome, and employed phylogenomics and reverse genetics in *Arabidopsis*, a model system that diverged from *Sphagnum* approximately 380 million years ago. Our analysis identified a specific gene encoding the proteasome-associated protein ECM29 as a critical factor in microbially conferred thermotolerance. Biochemical assays demonstrated that ECM29 inhibits *Arabidopsis* 26S proteasome activity by altering binding preference among the proteasome subunits in response to the bacterial strain *Variovorax* sp. CF313. This inhibition triggers the induction of heat-responsive genes, priming the plant for enhanced heat acclimation. The successful validation of this mechanism in *Arabidopsis* suggests potential evolutionary conservation of ECM29 function across major plant lineages from bryophytes to angiosperms.

Our findings underscore the value of ecological genomics in elucidating fundamental plant-microbe interactions. This study highlights the potential for leveraging microbial partnerships to enhance thermotolerance, with broad implications for improving resilience in food crops and bioenergy feedstocks amid a warming climate.

## Results

### QTL analysis identifies genes associated with host-microbe associations

Our goal was to investigate how plant genetic variation impacts microbiome interactions and their combined ability to withstand heat stress. Using a microbiome transfer approach (17, 22), we collected microbial communities sourced from *Sphagnum angustifolium* peat moss within the SPRUCE whole ecosystem warming experiment (29). Plants, and their associated microbiomes, were isolated from the warmest enclosure (+9°C above ambient temperatures) and ambient experimental enclosures. To infer genetic loci that cause variation in the ability of *Sphagnum* to receive microbially provided thermotolerance, we used clones from a recently developed F1-haploid pedigree population (30) co-cultured with either the warming or ambient sourced microbiomes using previously described methods (22). Log-transformed relative growth rate (hereafter ‘growth’, defined as the occupied area within each imaging well from time zero) was used as the phenotype in each experimental treatment. Growth benefit was calculated as the difference in growth per treatment relative to the control.

Within temperature conditions, microbiome sources provided distinct growth benefits (ambient chamber (21 °C); Wilcoxon; p=0.0002, warm chamber (30 °C); Wilcoxon; p=0.045). The greatest growth benefit occurred in plants grown in the warm chamber with warm-sourced microbiomes (+0.0014 mm^2^ d^-1^), followed by ambient source microbiomes (+0.0011 mm^2^ d^-1^) (Fig. 1A). The warm-sourced microbiome advantage showed a broad range of growth effects across individual genotypes of the pedigree within warming conditions (Fig. 1A). To validate the observed variation within the pedigree, genotypes exhibiting the highest (n=3) and lowest (n=3) growth benefits were independently co-cultured with either warming- or ambient-sourced microbiomes (6 replicates each). Under warming conditions, significant differences in growth benefit were observed between highest and lowest benefit genotypes as identified in the prior experiment (Wilcoxon; p<0.001). The greatest growth benefit occurred in the predicted high-benefit genotypes co-cultured with warming-sourced microbiomes (+0.86 mm^2^ d^-1^ compared to control) (Fig. 1B) suggesting host genetic variation plays a critical role in receiving thermal benefits from the microbiome.

**Figure 1.**
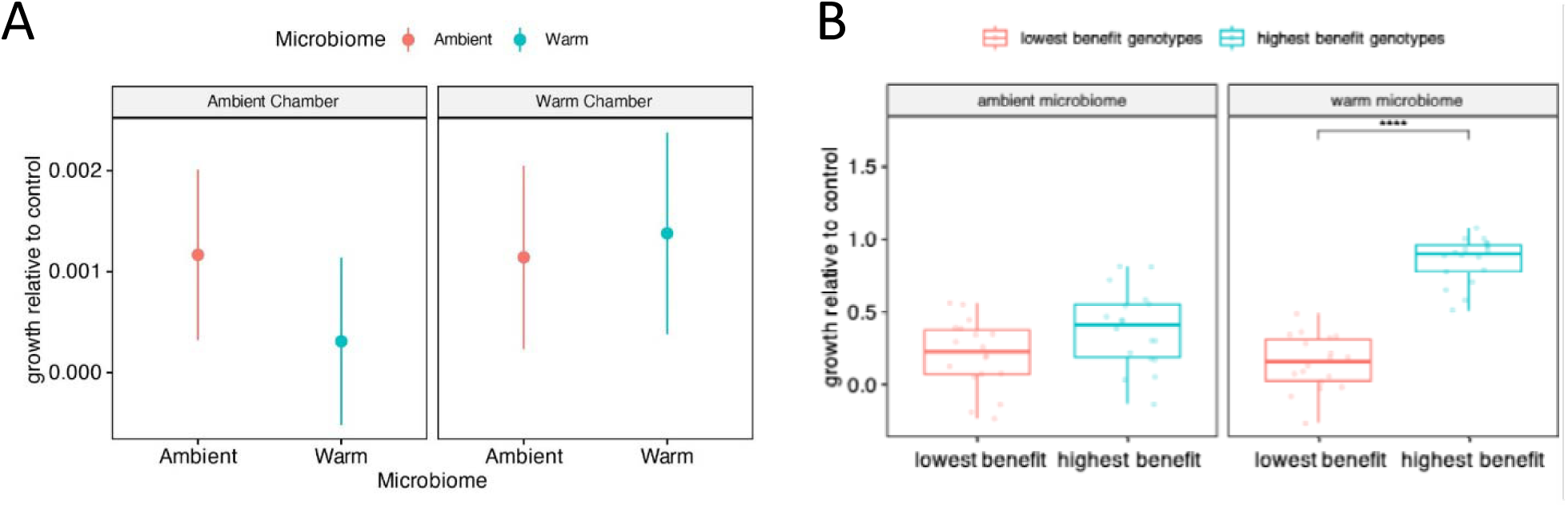
Microbiome benefit. Average growth rate of moss under ambient or warming conditions with whole microbial population **(A)** and extreme genotypes **(B)**. Error bars represent standard error of the mean of n = 6.

### *Variovorax* is involved in enhancing host plant thermotolerance

To identify specific microbial taxa that contribute to plant thermotolerance in controlled laboratory conditions, we aligned microbial 16S rDNA sequence reads from our previous *Sphagnum* microbiome transfer study (22) with 16S reads from our plant-microbe culture collection (23). Among the overlapping taxa, *Variovorax* emerged as a prominent candidate, showing strong enrichment in samples exhibiting microbially mediated thermotolerance. Its highest relative abundance (2.1-2.5%) was observed in elevated temperature conditions when plants received microbiomes preconditioned to elevated temperatures; a nearly 5-fold increase compared to plants that received ambient-conditioned microbiomes under the same heat stress (0.2-0.4%) (Fig. S1A).To experimentally validate the role of *Variovorax* in promoting thermotolerance, we performed single-strain inoculation experiments using two strains: *Variovorax* sp. CF313 (isolated from *Populus* trees), and *Variovorax* sp. IAA (isolated from *Sphagnum* within the SPRUCE experiment). Under elevated temperatures, uninoculated control plants exhibited significantly reduced growth (0.59 ± 0.39 mm² day⁻¹) compared to ambient conditions (1.62 ± 0.43 mm² day⁻¹). In contrast, plants inoculated with either *Variovorax* strain maintained growth under elevated temperatures (CF313: 1.21 ± 0.61; IAA: 1.71 ± 0.47 mm² day⁻¹) (Fig. S1B).

### ECM29 contributes to bacterially provided host plant thermotolerance

The QTL analysis identified a genomic interval on chromosome 13 containing seven gene models (Fig. 2; Table S1). To assess their role in microbially conferred thermotolerance, we used *Arabidopsis* as a heterologous genetic system due to the inability to genetically transform *Sphagnum*. The gene model Sphfalx13G087500 was excluded due to a lack of a homolog in *Arabidopsis*. Building on insights into key beneficial microbial taxa, which were further validated in *Arabidopsis* (24), we examined the interactions between *Arabidopsis* mutant lines and *Variovorax* sp. CF313. Of the six genes tested, only the null-mutant for gene model 5 (Sphfalx13G087900) displayed a phenotype indicative of a loss of microbially conferred thermotolerance. This gene shares homology with *ecm29* in *Arabidopsis* (AT2G26780), which encoded a proteasome-associated protein with a 71% amino acid similarity. ECM29 is a highly conserved protein, typically represented by a single gene family found across diverse plant species (Fig. S2) and contributes to the regulation and stabilization of the 26S proteasome complex that is essential for protein degradation in eukaryotic cells.(31, 32). The absence of observable phenotypes for the other gene models may be due to functional redundancy within multigene families. For instance, AT2G17280, a homolog of Sphfalx13G087700 that encodes phosphoglycerate mutase, has been reported to respond to fungal elicitors and nematodes (33, 34). Another candidate, Sphfalx13G088000, encodes a Rho family GTPase and warrants further investigation due to its role in cytoskeletal rearrangement during symbiotic interactions (35, 36). Given the bioassay results, its single-copy status in most plant species (including *Sphagnum* and *Arabidopsis)*, and its role in proteasomal regulation, we employed *Arabidopsis ecm29* mutants to test whether ECM29-mediated proteasomal control contributes to microbial-provided thermotolerance.

**Figure 2.**
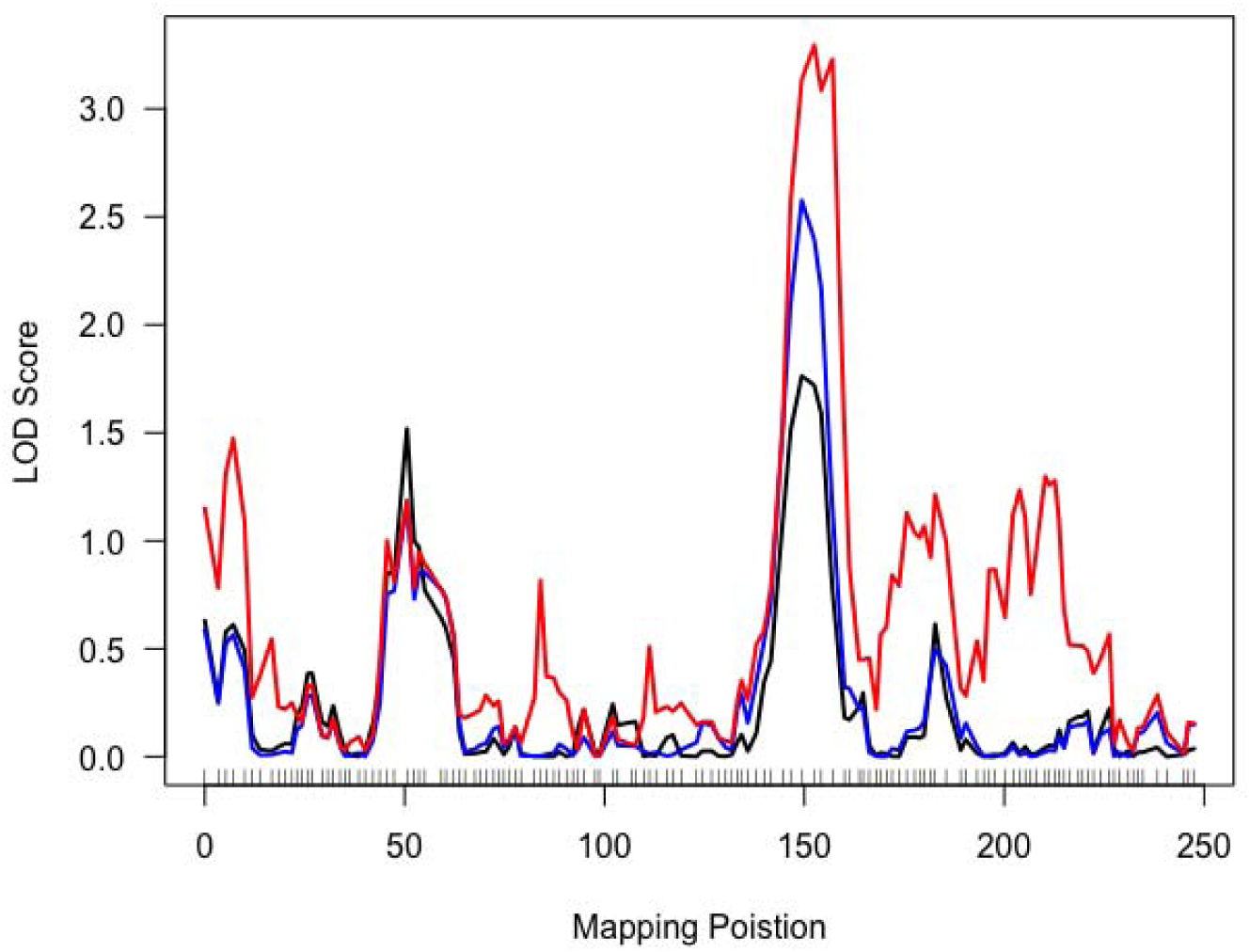
QTL mapping of high temperature growth differences from microbiome co-cultures. One QTL peak was detected on chr.D13 LOD scores, conditional on other QTL in a multiple QTL model, are presented. The QTL is dependent on sex and autosomal parental allele (blue, A allele; red, B allele).

The *Arabidopsis ecm29* gene (AT2G26780) spans approximately 13 kb and consists of 35 exons. In this study, we used two independent mutant lines: *ecm29-1* (SAIL_759_B04) and *ecm29-3* (SALK_137715), which harbor T-DNA insertions in the 15^th^ and 28^th^ exons, respectively (Fig. 3A). Homozygous mutants were identified by PCR using allele-specific primers (Fig.3B). *ecm29* transcripts were undetectable in *ecm29-1* plants and were present at very low abundance levels in *ecm29-3* plants relative to wild-type plants (Fig.3C). When exposed to non-lethal thermopriming (37°C for 90 min) prior to heat shock (45°C), both mutant and wild-type plants exhibited enhanced thermotolerance, indicating that ECM29 is not required for innate heat priming (Fig. 3D). However, unlike wild-type plants, *ecm29-1* and *ecm29-3* mutants failed to acquire bacterium-mediated thermotolerance following inoculation with *Variovorax* sp. CF313, suggesting that ECM29 is essential for microbe-enabled thermal resilience (Fig. 3D). Transgenic *Arabidopsis* lines expressing *ECM29* under the constitutive AtUBQ10 promoter (overexpression lines, OE) or its native promoter (complementation lines, Native) in the *ecm29-1* background (Fig. S3A) were generated to further characterize *ecm29* function. In OE lines, *ecm29* transcript levels were elevated 4- to 10-fold relative to wild-type, while complementation lines restored expression levels to near wild-type levels (Fig. S3B). In both cases, bacterially conferred thermotolerance was restored (Fig. S3C), confirming that ECM29 is necessary for the beneficial effects of *Variovorax* under thermal stress.

**Figure 3.**
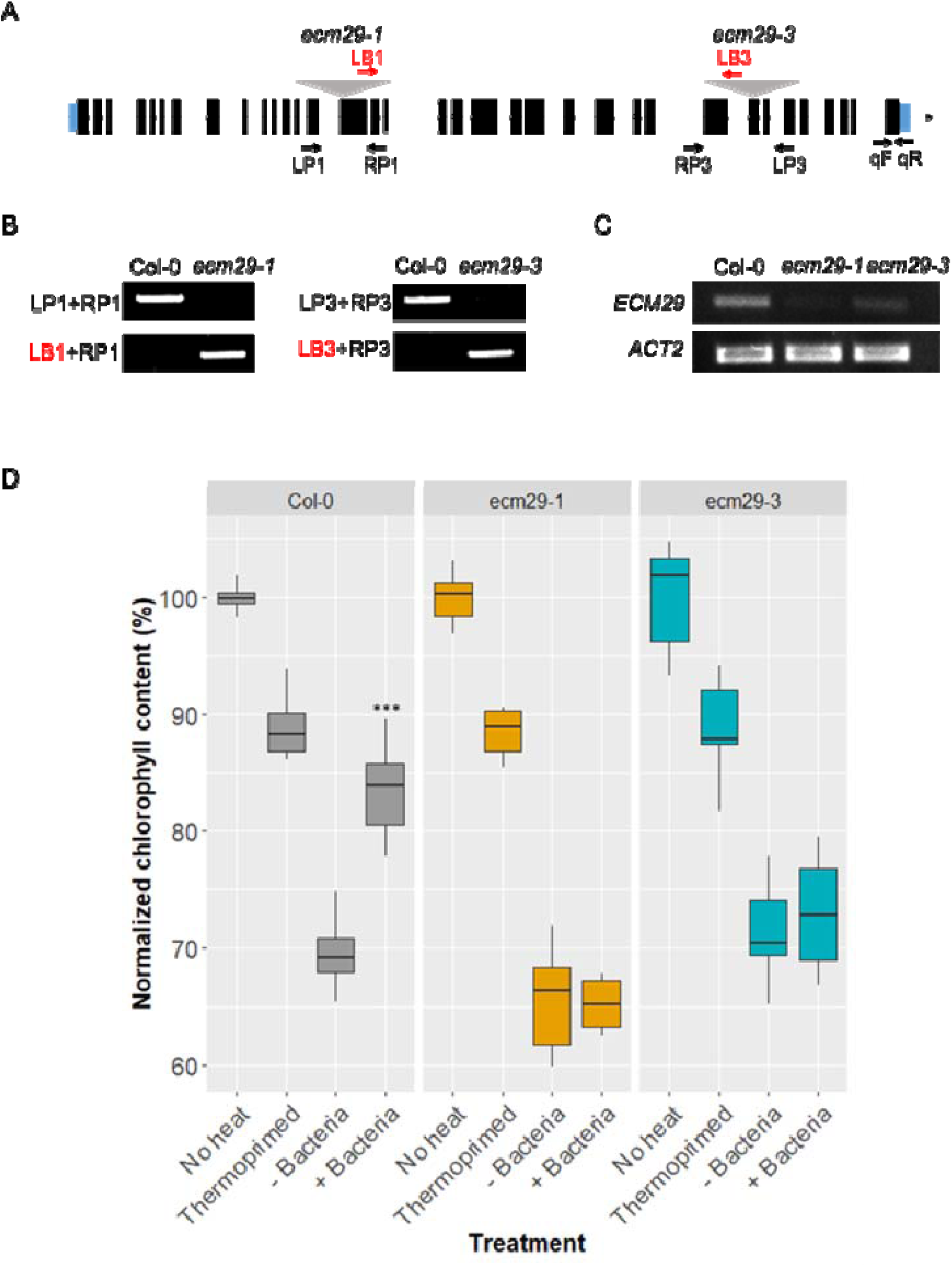
*ecm29* T-DNA insertional mutants are compromised in bacterially provided thermotolerance. **(A)** Schematic illustrates T-DNA locations (gray triangles) in the 15th and 28th exons (black bar) in ECM29 for ecm29-1 and ecm29-3, respectively. Blue boxes indicate 5′- and 3′-untranslated regions. Arrows indicate the positions of primers. **(B)** Homozygosity of the T-DNA insertion in ecm29-1 and ecm29-3 was determined by PCR analysis of the ECM29 genomic fragment using left primers (LP1 and LP3 for ecm29-1 and ecm29-3, respectively) and right primers (RP1 and RP3 for ecm29-1 and ecm29-3, respectively), and the T-DNA insertion using a T-DNA-specific left border primers (LB1 and LB3 for ecm29-1 and ecm29-3, respectively) with RPs. Col-0 is the wild type. **(C)** ECM29 expression level in wild type, ecm29-1, and ecm29-3 plants was determined by RT-PCR analysis with primers (qF and qR, ECM29 specific) and ACTIN2 (ACT2, reference standard). **(D)** Total chlorophyll content was measured from wild type, ecm29-1, and ecm29-3 plants without heat shock or with heat shock after thermopriming or with/without bacterial inoculation. Box plots display the 25th – 75th percentiles with the median (horizontal line) (n = 5 - 17 wells with ca. 10 seedlings each). ***, *P*<0.001.

### ECM29 dynamically interacts with 26S proteasome subunits in the presence of bacteria

ECM29 has been reported to bind to multiple 26S proteasome subunits in human and yeast systems, acting as a regulatory hub that modulates proteasomal assembly, disassembly, and activity (31, 32, 37, 38). To investigate ECM29–proteasome interactions in plants and assess how these interactions change in response to *Variovorax*, we used *Arabidopsis* seedlings expressing ECM29 fused to a 3×HA-YFP tag for immunoprecipitation (IP) followed by liquid chromatography–tandem mass spectrometry (LC-MS/MS).

As in human and yeast systems, multiple Arabidopsis core proteasome subunits and associated factors were identified as ECM29 interactors under both uninoculated and co-culture conditions (Dataset S1). These included key components of the 20S core particle (CP) and the 19S regulatory particle (RP) (Table 1). Notably, ECM29 was more likely to be associated with the 19S RP, lid subunits, (e.g., RPN5) and base subunits (e.g., RPN1, RPT2, RPT3, RPT4, RPT5, and RPT6) regardless of bacterial presence. Under inoculated conditions, however, ECM29 additionally co-precipitated with several additional 19S RP subunits (e.g., RPN6, RPN8, RPN12, and RPT1) and 20S CP subunits (PAC and PAD), suggesting that ECM29 dynamically alters its binding partners in response to bacteria. To substantiate these findings, we performed Co-IP experiments using anti-bodies raised against PAC1, a 20S CP subunit, and RPT1, a 19S RP subunit. Co-IP revealed strong signals for PAC and RPT1 in inoculated samples, despite a weaker GFP signal, confirming that these subunits preferentially bind to ECM29 in the presence of *Variovorax* sp. CF313 (Fig.4A). These results underscore the dynamic interaction of ECM29 with the 26S proteasome in response to bacterial cues.

**Figure 4.**
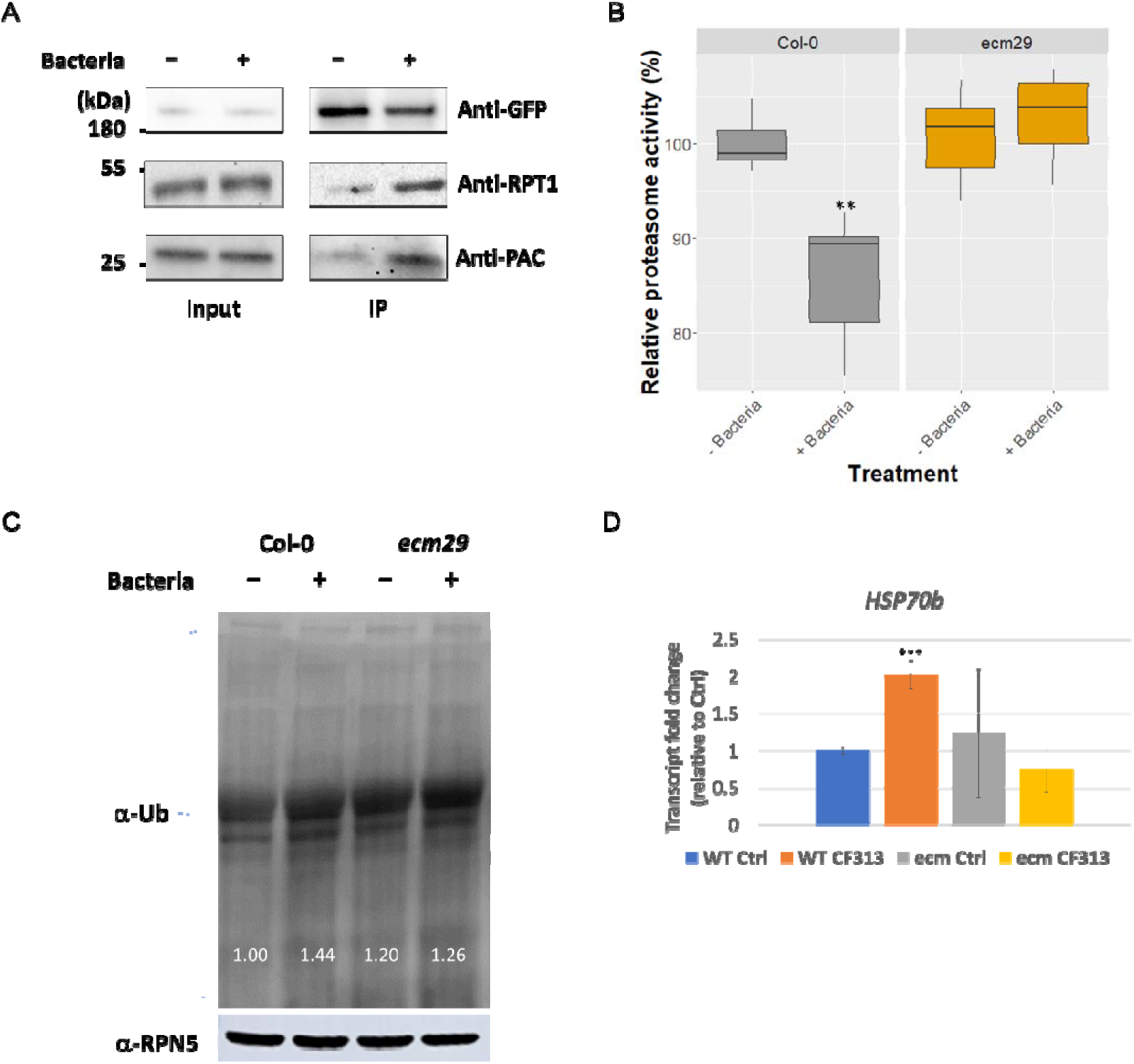
ECM29-mediated regulation of proteasome activity upon the presence of bacteria which changes interacting partners. **(A)** ECM29-HA-YFP was pulled-down using Pierce^TM^ anti-HA magnetic beads. ECM29-HA-YFP, RPT1, and PAC from the input and pulldown fractions were detected by immunoblots using anti-GFP, anti-RPT1, and anti-PAC antibodies, respectively. The results show that ECM29 preferentially interacts with the 26S proteasome subunits, RPT1 and PAC, in the presence of bacteria. **(B)** Bacteria-induced inhibition of proteasome activity was observed in wild-type plants, but not in *ecm29* mutant plants. Box plots display the 25th – 75th percentiles with the median (horizontal line) (n = 6). **(C)** Western blot of polyubiquitinated (Ub) proteins and **(D)** relative gene expression levels of *HSP70b.* **, *P*<0.05; ***, *P*<0.001.

Given the bacterial-induced shift in ECM29 binding partners, we hypothesized that plant-bacterial interactions regulate proteasome activity via ECM29. To test this, we measured proteasome activity in wild-type and *ecm29* mutant plants, with and without bacterial inoculation. In wild-type plants, *Variovorax* sp. CF313 presence suppressed 26S proteasome activity, whereas no inhibition was observed in *ecm29* mutants (Fig. 4B), indicating that ECM29 is required for bacterial-mediated suppression of proteasome activity.

Proteasome dysfunction leads to the accumulation of proteins marked for degradation, triggering the induction of chaperones such as heat shock proteins (HSPs) to mitigate proteotoxic stress. To determine if bacteria-induced proteasome inhibition triggers a similar response, we assessed the ubiquitination status of host proteins across co-culture conditions. Consistent with prior findings, inoculated wild-type plants exhibited a ∼44% increase in ubiquitinated proteins relative to uninoculated controls (Fig.4C). This accumulation is consistent with known proteasome inhibition by compounds like MG132 and Syringolin A, which are known to induce HSP expression (39, 40). Accordingly, we observed significant induction of *HSP70b* transcripts in inoculated wild-type plants (Fig. 4D). Elevated basal levels of HSP70 are associated with enhanced thermotolerance by enabling plants to better manage heat-induced proteotoxicity (41–43).

## Discussion

Microbes can enhance plant thermotolerance through various mechanisms, including the induction of heat-responsive genes, production of protective metabolites, and modulation of nutrient allocation and composition (18–20). In this study, we combined a microbiome transfer approach with QTL mapping using an ecologically relevant *Sphagnum* pedigree to identify a seven-gene interval associated with microbially conferred thermotolerance. Using a heterologous *Arabidopsis* system, we investigated genes within the QTL interval and identified *ecm29* (Sphfalx13G087900; AT2G26780) as essential, as mutant lines failed to exhibit microbially conferred thermotolerance. Further, we showed that a common *Variovorax* strain, frequently found in plant microbiomes exposed to warming conditions, modulates ECM29 to influence the host ubiquitin-proteasome system (UPS), a key regulator of protein homeostasis, abiotic stress signaling, and defense response.

Plant-microbe partnerships present a promising strategy to enhance acclimation and resilience to environmental stress. Microbiomes adapted to extreme environments, capable of rapidly adjusting to fluctuating conditions, can confer stress tolerance to their plant hosts by transferring stress tolerance traits (3, 44). For instance, Allsup et al. (2023) demonstrated that tree seedlings inoculated with microbial communities from cold or dry habitats improved survival and tolerance to drought, heat, and cold stress (17). Similarly, Carrell et al. (2022) used a whole-peatland warming study to test the habitat-adapted microbiome concept, transferring microbiomes from field samples to axenic laboratory-grown plants, which demonstrated enhanced resilience to heat stress (22). Despite these documented benefits, the molecular genetic mechanisms by which plants harness microbial benefits to abiotic stress remain largely unexplored. Our study provides a novel mechanistic understanding of how host genetic factors interact with microbial partners to enhance thermotolerance.

A recent study by Shekhawat et al. (2021) offers insights into how microbial symbionts influence host thermotolerance (20). The researchers demonstrated that endophytic bacteria can mediate host epigenetic modifications, regulating the expression of heat stress memory genes.

Specifically, *Enterobacter* sp. SA187 induced constitutive *H3K4me3* modifications at the promoter regions of heat stress memory genes (e.g., *hsp18.2* and *apx2*) via the ethylene pathway, leading to enhanced plant thermotolerance (20). In contrast, our study found no significant transcriptional changes in *hsp17*.6 and *apx2* in response to *Variovorax* sp. CF313 (Fig. S4), suggesting alternative mechanisms. Instead, our results reveal that *Variovorax* enhances plant thermotolerance by modulating the host ubiquitin-proteasome system (UPS). This highlights the diversity of mechanisms that microbes utilize to confer similar phenotypic benefits to their host plants and suggests that multiple evolutionary strategies have emerged to mediate plant-microbe thermotolerance.

The UPS is critical for plant development and responses to environmental stress, especially in sessile organisms like plants that cannot escape fluctuating conditions (45). This system is particularly relevant to thermotolerance as it regulates the degradation of misfolded or damaged proteins, preventing protein aggregation under heat stress and maintaining cellular homeostasis. Target proteins are ubiquitinated through a sequential enzymatic cascade involving E1 ubiquitin-activating enzymes, E2 ubiquitin-conjugating enzymes, and E3 ubiquitin ligases, after which the ubiquitinated proteins are degraded by the 26S proteasome. By selectively degrading key proteins such as transcription factors, the UPS modulates acclimation, stress response and recovery (46). The UPS also plays a pivotal role in plant immunity, from pathogen recognition to the execution of the immune response (47). The involvement of the UPS in plant responses to both biotic and abiotic stressors suggests that it serves as a central regulatory hub orchestrating various stress-related mechanisms.

Pathogenic bacteria have evolved strategies to manipulate the plant UPS, suppressing host defenses and promoting disease (47, 48). For example, bacterial effectors can manipulate the plant UPS pathway by promoting ubiquitination of immunity-associated proteins, leading to the suppression of defense responses (49); mimic plant F-box proteins to hijack the UPS and facilitate disease (50); or directly inhibit proteasome activity (51, 52). While the role of the UPS in plant stress responses is well established (53), whether beneficial microbes also modulate the plant proteasome during symbiosis remains unclear. The 26S proteasome is the key component of the UPS, responsible for degrading 80-90% of cytosolic and nuclear proteins (54). It consists of a 20S core particle (CP) capped by one or two 19S regulatory particles (RP) (55, 56). The CP contains two outer α-subunit rings that form a gate and two inner β-subunit rings where proteolysis occurs. The 19S RP is responsible for substrate recognition, de-ubiquitination, unfolding, and translocation into the proteolytic 20S CP. Its base contains six ATPase subunits (RPT1-6), which are essential for substrate unfolding and opening the α-ring gate of the 20S CP (57).

We found that *Variovorax* sp. CF313 reduces host proteasomal activity through ECM29, which preferentially binds to RPT1 and leads to the accumulation of ubiquitinated proteins. This, in turn, induces elevated expression levels of *hsp70b*, a heat-responsive chaperone crucial for maintaining protein homeostasis, even in the absence of heat stress. This primes the plant for subsequent heat shock (58, 59). In mammalian cells, proteasome inhibition similarly induces HSPs and enhances thermotolerance (40, 60, 61). ECM29 has been primarily studied in yeast and mammalian cells, where it regulates proteasome assembly and activity. Proposed mechanisms include stabilizing the proteasome complex by tethering RP and CP (62) and facilitating dissociation under oxidative stress (63), and repressing proteasome function by inhibiting RPT ATPase activity in RPTs and regulation of the CP gate (64), which aligns with our findings. Our findings expand the functional characterization of ECM29 to plants and identify its novel role in mediating plant-microbe interactions. The observed modulation of proteasome activity suggests that ECM29-dependent proteasome regulation may represent an evolutionarily conserved mechanism for plant response to microbial signals.

Using proteomics and co-immunoprecipitation, we show that ECM29 interacts with different subunits of the 26S proteasome complex in response to *Variovorax* sp. CF313, a growth-promoting bacterium enriched in our field warming microbiomes (Fig. S2A). Regardless of bacterial presence, ECM29 binds to 19S RP subunits including RPT4 and RPT5 likely through its N-terminal region. This is consistent with previous studies in human cells (63), where cross-linking mass spectrometry identified ECM29 interactions with RPT1, RPT3, RPT4, RPT5, RPN1, and RPN10 (55). Wang et al. (2017) proposed that ECM29 is recruited to the 19S RP in response to oxidative stress, promoting free 20S CP activity (63). A key finding from the current study is that, in the presence of *Variovorax,* ECM29 specifically binds to RPT1. The C-terminal tails of RPT2, RPT3, RPT5 contain a conserved HbYX motif (hydrophobic-tyrosine-variable C-terminal residue), which insert into α-subunit pockets to regulate 20S CP gate, functioning as a “key-in-lock” mechanism (65). Cryo-electron microscopy studies have shown that the HbYX motifs of RPT2, RPT3, and RPT5 dock into the α pockets in both closed- and open-gate states, but sufficiently to trigger gate-opening (66, 67). Although RPT1 has only a partial HbYX motif, it plays a key role in opening the 20S CP gate by repositioning structural elements of the α-subunits through its C-terminal tail (68). Our data show that ECM29 binding to RPT1 moderately inhibits 20S CP gate opening, resulting in a 10% to 20% reduction in proteasome activity (Fig. 4B). This interaction reveals a nuanced regulatory mechanism by which *Variovorax* influences host proteasome dynamics to enhance plant thermotolerance.

Taken together, our conceptual model (Fig. 5) suggests that yet-to-be-identified bacterial factors alter ECM29 binding specificity within the 26S proteasome, potentially through conformational alterations or the displacement of a repressor at the binding site. This modification enables ECM29 to bind the RPT1 tail, thereby inhibiting the stability of 20S CP gate opening and reducing proteasome activity. The resulting accumulation of ubiquitinated proteins induces *hsp70* expression, priming the plant for improved thermotolerance. While our study has identified the host genetic component of this interaction, future work should focus on identifying the bacterial signals that initiate this process. Potential candidates include secreted proteins, lipopolysaccharides, or small molecules that could interact directly with ECM29 or indirectly through other host proteins. While this thermotolerance may partly reflect immune suppression during symbiosis, it likely contributes to broader organismal and ecosystem resilience. In this study, *Sphagnum*—a key ecosystem engineer that is highly sensitive to warming (69, 70)— benefited from microbial conferred thermotolerance. Given its central role global carbon storage, these findings underscore the importance of enhancing plant and ecosystems resilience to climate stressors.

**Figure 5.**
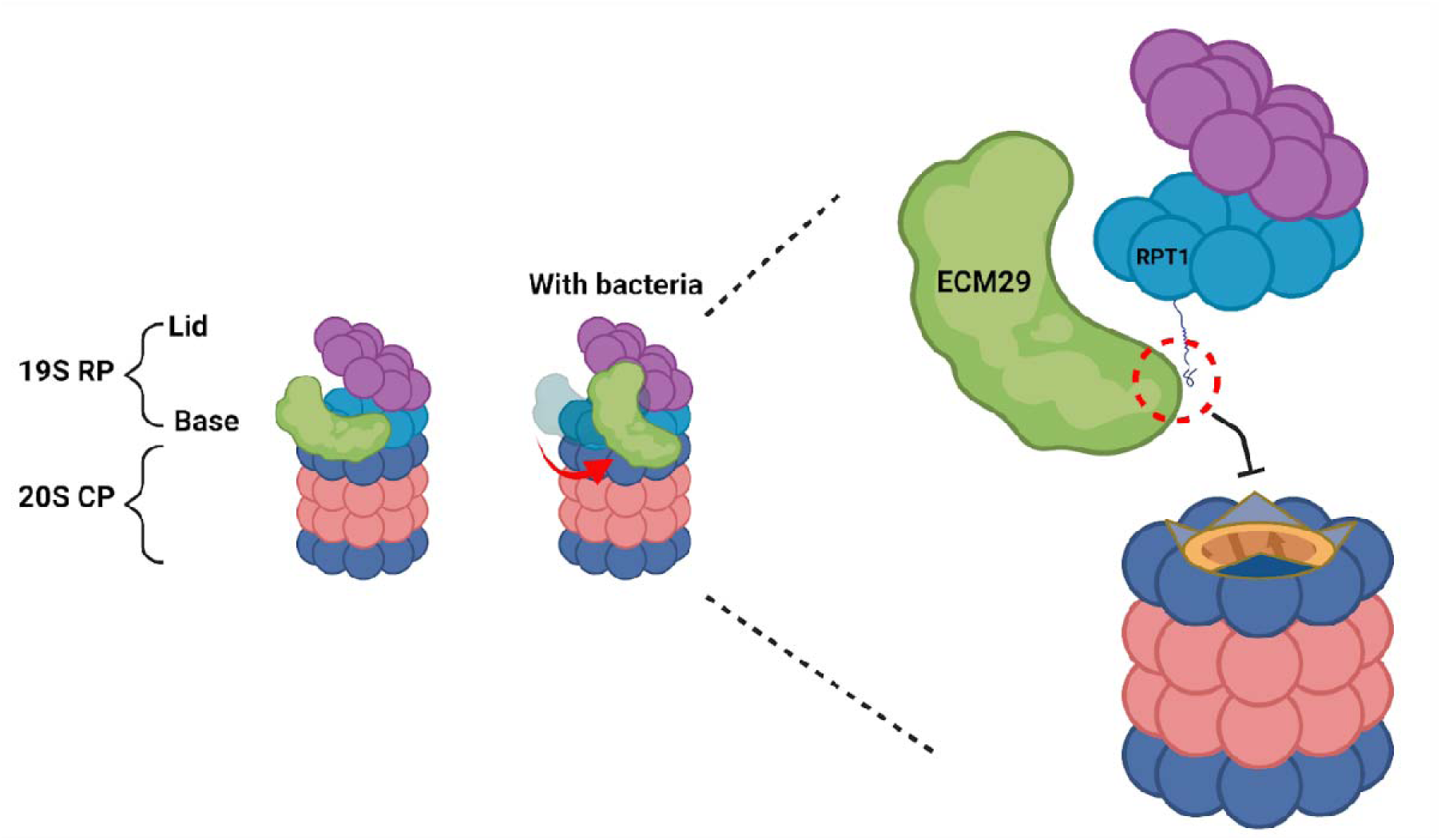
Conceptual model of ECM29-dependent regulation of the 26S proteasome upon interaction with *Variovorax*. ECM29 interacts with multiple subunits of both the 19S and 20S proteasome, modulating the activity and stability of the 26S proteasome. In the presence of the beneficial bacteria *Variovorax*, ECM29 specifically binds to RPT1, leading to moderate inhibition of the 20S proteasome gate opening and a consequent reduction in proteasome activity.

The conservation of ECM29 function between *Sphagnum* and *Arabidopsis*, plants that diverged approximately 380 million years ago, suggests that this mechanism may be widespread across the plant kingdom. This conservation provides an opportunity to engineer enhanced thermotolerance in crops by optimizing microbiome composition or enhancing the plant’s ability to receive microbial benefits. As climate change continues to increase heat stress frequency and intensity, such approaches could help maintain agricultural productivity and ecosystem function.

Our findings suggest that climate resilience is not solely determined by innate stress tolerance at the organismal level, but is deeply influenced by species interactions and including intricate plant-microbial interactions. This perspective shifts our understanding of climate adaptation from a purely genetic trait to one that incorporates the extended phenotype conferred by associated microbes. Future studies should investigate (1) the identifying bacterial signals that influence ECM29*-*proteasome interactions, (2) assessing conservation of these mechanisms across plant taxa and other plant-microbe interactions, (3) leveraging this knowledge to improve restoration and resilience, and 4) exploring whether similar mechanisms mediate other forms of microbially-conferred stress tolerance, such as drought or salinity tolerance, to reveal common mechanisms underlying beneficial plant-microbe interactions.

In conclusion, this study integrates ecological and molecular approaches to elucidate a novel mechanism whereby beneficial microbes enhance plant thermotolerance. We demonstrate that *Variovorax* modulates ECM29-dependent proteasome activity, priming plants for heat stress without prior thermal exposure. This finding not only advances our fundamental understanding of plant-microbe interactions but also provides potential strategies for enhancing ecosystem and agricultural resilience.

## Materials and Methods

### Conditioned microbiome transfer experiment

Field-conditioned microbiomes were obtained from Sphagnum material collected at the Spruce and Peatland Responses Under Changing Environments (SPRUCE) experiment and applied to the axenic Sphagnum pedigree following methods described in “Habitat-adapted microbial communities mediate Sphagnum peatmoss resilience to warming”. Briefly, we collected 100 g of living green stems of hollow-dwelling Sphagnum angustifolium from ambient +0°C (ambient) and ambient +9°C (elevated) plots. Samples were placed in individual sterile bags and shipped to the laboratory on blue ice. Microbiomes were isolated by dicing 100 g of tissue with a sterile razor blade and pulverizing it in PBS with a mortar and pestle. The resulting suspension was filtered, centrifuged to pellet the microbes, and resuspended in 500 ml BG11-N medium (pH 5.5). Each suspension was applied to clones from a recently developed F1-haploid pedigree population of Sphagnum. A single capitulum of each genotype was added to each well of a 12-well plate and inoculated with 2 ml of ambient-microbiome, warm-microbiome, or sterile media.

Sealed plates were placed into growth chambers with 350 μmol m^-2^ s^-1^ of photosynthetically active radiation (PAR), 12 h: 12 h, light: dark cycle, programmed to either ambient (16.75°C average temperature) or elevated (25.75°C average temperature) field plot temperatures (Table S2. for full temperature settings). To measure growth, we collected images from the top of each plate weekly and measured surface area using ImageJ software. The change in surface area was determined as a proxy for growth. Log-transformed relative growth rate (defined as the occupied area within each imaging well from time zero) was used as the phenotype in each experimental treatment. Growth benefit was calculated as the difference in growth per treatment relative to the control. We compared the growth rate benefits across microbiome sources and growth chambers using Wilcoxon tests.

To confirm growth benefit results, we selected genotypes with the most (n=3) and least (n=3) growth benefits from the pedigree experiment. These genotypes were co-cultured with warming or ambient sourced microbiomes (6 replicates each). Under warming conditions, we compared growth benefits between the highest and least benefit genotypes using a Wilcoxon test.

### QTL analysis

Quantitative loci mapping was performed in R/qtl (v.1.50) using the Haley–Knott regression method on hidden Markov model-calculated genotype probabilities as described previously (71). Briefly, one-way and multiple QTL model scans were conducted on log-transformed phenotypes to correct for right-skewed distributions. In all QTL scans, sex was treated as a covariate as determined both by markers extracted from the genetic map for chr. 20, as well as the ratio of reads mapped to the shared region among chr. 20 and Scaffold9707 as described previously (71). Any sample where the marker data or mapping ratio were ambiguous was assigned ‘NA’. To account for plate-specific variation, week 2 measurements were normalized by subtracting the plate-wise mean and adding one. These normalized values were used to compute adjusted week 2 and week 4 values. Relative growth rate (RGR) was calculated as the change in log-transformed values over the two-week interval using normalized measurements. To determine the significance thresholds for each QTL, 1,000 permutations were performed. To estimate the effects of predicted sex (male and female), genotype at each QTL locus (A or B) and treatment (ambient and warming) on log-transformed growth, we fit a univariate mixed linear model with all two- and three-way interactions, with a random effect of individuals to account for the same individual measured in two conditions. Sex-specific linear models were run with interactions, with goodness of fit compared between each. *S. angustifolium* genes within significance intervals are listed in Supplementary Table1.

### *Arabidopsis* genotypes and growth conditions

*Arabidopsis thaliana* genetic resources used in this study were wild type (Col-0) and two mutant lines *ecm29-1* (SAIL_759_B04) and *ecm29-3* (SALK_137715) obtained from the Arabidopsis Biological Resource Center at Ohio State University. Using *ecm29-*1, transgenic plants expressing AtUBQ10_pro_:ECM29-3×HA-YFP and AtECM29_pro_:ECM29 were generated by *Agrobacterium tumefaciens*-mediated floral dip method (72), and then single-copy homozygous T_3_ plants were used for further analysis. Surface sterilized seeds with 70% (v/v) ethanol and bleach solution 50% (v/v) bleach/0.05% (v/v) Tween-20] were stratified for 3 d at 4°C in the dark, and then planted on autoclaved PTFE mesh discs (0.025”× 0.005” opening, McMaster-Carr) placed onto medium plate 1x Murashige and Skoog (MS) basal medium with vitamins, 0.5 g/L MES, pH 5.7, and 0.7% agar], as described in (24). One-week-old seedlings on mesh discs were then transferred to 6-well plates (VWR) that contained 3 ml of 1x liquid MS medium in each well with or without bacteria. Plants were grown in a growth chamber (Precision^TM^ Plant Growth Chamber, Thermo Scientific) under a 12 h light cycle with 100 µmol photon m^-2^s^-1^ at 22°C.

### Bacteria preparation and plant inoculation

*Variovorax* spp. CF313 and IAA were maintained on R2A agar plates and cultured at 28°C for two days. A single bacterial colony from the agar plate was inoculated into 3 mL of R2A broth and cultured at 28°C with shaking at 250 rpm overnight, spun down, and the pellet was washed 3 times with liquid MS medium. The OD600 was determined using a BioTek microplate reader, and then diluted to a final OD600 of 0.01 in liquid MS medium before adding to a 6-well plate to inoculate seedlings.

### Thermotolerance analysis

Heat shock treatment followed by chlorophyll content measurement were performed according to previously established method (24). Briefly, 2-week-old seedlings with or without bacteria were subjected to heat shock for 14 min at 45°C, placed back to normal growth condition for recovery for 4 d, then harvested for extracting total chlorophyll in 100% DMSO. The concentrations of total chlorophyll (mg/g F.W.) was quantified using the following equation: 7.49*A*^665^ + 20.34*A*^648^.

### Co-immunoprecipitation and immunoblots

Transgenic seedlings (AtUBQ10_pro_:ECM29-3×HA-YFP) grown with or without bacteria (2-3 g) were harvested and ground using Sample Prep 2010 Geno/Grinder (SPEX Sample Pre, UK). Powder was homogenized in 6 mL extraction buffer 50 mM Tris-Cl pH 7.5, 100 mM NaCl, 1 mM EDTA, 0.1% NP40, 40 µM MG132, 40 µM MG115, 1× cOmplete^TM^ protease inhibitor cocktail (MilliporeSigma, Burlington, MA)], centrifuged at 20,000 × *g* for 20 min at 4°C, then crude extract was cleared by 0.45 µm polyethersulfone filter. Cleared extract was mixed with 100 µl equilibrated anti-HA affinity matrix (Roche) at 4°C for 2h, then the matrix was washed 4 times with 1ml extraction buffer. Bound proteins were eluted by boiling for 10 min in 50µl 2× SDS sample buffer and used for proteomic analysis and immunoblotting. Transgenic seedlings expressing 3×HA-YFP only were used as a negative control to exclude non-specific binding proteins.

For immunoblotting, cleared protein samples (20 µl) were separated on 4–15% Mini-PROTEAN® TGX™ Precast Protein Gel (Bio-Rad Laboratories) and then transferred to nitrocellulose membranes. The membranes were incubated with primary antibodies polyclonal anti-GFP antibodies (Thermo Fisher Scientific, A-11122) and polyclonal rabbit anti-PAC1 antibodies (Agrisera, AS19 4258)] at 1:3000 dilution, then incubated with horseradish peroxidase-conjugated goat anti-rabbit secondary antibodies (Agrisera, AS09 602). Signals were detected via chemiluminescence using a SuperSignal™ West Pico PLUS kit (Thermo Fisher Scientific, 34577). For detecting polyubiquitinated proteins, monoclonal mouse anti-ubiquitin antibodies (Santa Cruz, sc-8017) and goat anti-mouse antibodies (Licor 926-32210) for primary and secondary antibodies, respectively.

### Proteomic analysis

For each eluent (120 µL), 80 µL of lysis buffer (20% sodium dodecyl sulfate) containing 10mM diothiothreitol was added to denature proteins. Samples were vortexed and incubated for 10 minutes at 90°C with constant shaking. The denatured protein sample was vortexed and centrifuged at 21,000xg for 10 minutes. Denatured proteins were alkylated with 30mM iodoacetamide and incubated in the dark for 15 minutes to prevent the reformation of disulfide bonds. Sera-Mag beads were added (2 µL) and protein aggregation capture was performed. Acetonitrile (ACN) was added to each sample to reach a final concentration of 70%. Samples were vortexed, allowed to settle for 10 minutes, vortexed, and allowed to settle for a final 10 minutes. Tubes were placed on a magnetic rack and the supernatant was removed using a vacuum system. Samples were washed with 1mL ACN and 1 mL 70% ethanol while on the magnetic rack. Samples were removed from the magnetic rack and beads were resuspended in 100mM ammonium bicarbonate (ABC). Proteins were digested using two aliquots of sequencing grade trypsin (1 µg) overnight, followed by 3h at 37°C with constant shaking. The samples were placed on a magnetic rack and adjusted to 0.5% formic acid (FA). A AcroPrep Advance 96 well 10 KDa omega filter plate (Pall Corporation) was prepped by adding 100 µL of ABC to each well and centrifuged at 1500xg for 10 minutes. Tryptic peptide samples were then arrayed in the filter plate and centrifuged at 1500xg for 30 minutes. Samples were freeze dried and then resuspended in 20ul aqueous solvent (5% ACN and 0.1% FA). Tryptic peptide concentrations were assessed using the Nanodrop One spectrophotometer. Peptide mixtures were analyzed using one-dimensional liquid chromatography on an Ultimate 3000 RSLCnano system (Thermo-Fisher Scientific) coupled with a Q Exactive Plus mass spectrometer (Thermo-Fisher Scientific). For each sample, a 2-μg inject of each sample was flowed across an in-house-built reversed-phase (RP) C18 trap column (5 µm by 150 µm by 50 mm) using an aqueous solvent (5% ACN and 0.1% FA) for 10 min. The trapped peptides were separated by a 90-min linear organic gradient (300 nL/min flow rate) from 2% organic solvent (80% ACN and 0.1% FA) to 30% organic solvent to separate peptides across an in-house-pulled nanospray analytical column (75 µm by 350 mm) packed with C18 Kinetex RP C18 resin (1.7 µm) (Phenomenex). All MS data were acquired with Thermo Xcalibur (version 4.2.47) using the top N method as previously described (73).

All MS raw data files were analyzed using the Proteome Discoverer software (Thermo-Fisher Scientific, version 2.1) (74). Each MS raw data file was processed by the SEQUEST HT database search algorithm (75) and confidence in peptide-to-spectrum (PSM) matching was evaluated by Percolator (76). The protein reference databases used for this study were the following: Araport11, *Variovorax* sp. CF313, and the ECM29-3×HA-YFP construct. Peptide and PSMs were considered identified at q < 0.01 and proteins were required to have at least one unique peptide sequence. Proteins that were absent in the negative control construct (3×HA-YFP) but detected in the samples expressing ECM29-3×HA-YFP were considered candidate interacting proteins.

### Proteasome activity assay

Proteasome activity assay was performed as described in Üstün et al. (2017) with a slight modification (77). To extract total protein, two-week-old seedlings with or without bacteria (50-100 mg) were harvested and ground to a fine powder in liquid nitrogen, then homogenized in 200 µl extraction buffer 50 mM HEPES-KOH (pH 7.2), 2 mM ATP, 2 mM DTT, 0.25 M sucrose]. After centrifugation, the supernatant was used to measure the protein content, and then the concentration of the total protein was adjusted to 1 mg/ml with an extraction buffer. To measure proteasome activity, 8 µg of total protein was added to a black walled 96-well plate and subsequently mixed with 150 µl assay buffer 100 mM HEPES-KOH (pH 7.8), 2 mM ATP, 5 mM MgCl_2_, 10 mM KCl]. After incubating for 15 min at 30°C, proteasome substrate labeled with a fluorescent 7-amino-4-methylcoumarin (AMC) was added to the final concentration of 100 µM: succinyl-Leu-Leu-Val-Tyr-AMC (MilliporeSigma, S6510), Z-Ala-Arg-Arg-AMC (MilliporeSigma, 539149), and Z-Leu-Leu-Glu-AMC (MilliporeSigma, C0483), then the release of free AMC was measured using a BioTek microplate reader at an excitation wavelength of 360 nm and an emission wavelength of 460 nm every 1.5 min for 2 hrs.

## Acknowledgments

This research was sponsored by the Genomic Science Program, U.S. Department of Energy, Office of Science, Biological and Environmental Research, as part of the Plant Microbe Interfaces Scientific Focus Area (http://pmi.ornl.gov). Oak Ridge National Laboratory is managed by UT-Battelle, LLC, for the U.S. Department of Energy under contract DE-AC05-00OR22725. The work (proposal no. 10.46936/10.25585/60001200 and 10.46936/10.25585/60001030) (A.J.S.) conducted by the US Department of Energy (DOE) Joint Genome Institute (https://ror.org/04xm1d337), a DOE Office of Science User Facility, is supported under contract no. DE-AC02-05CH11231.

## Notes

### Competing Interest Statement

The authors have declared no competing interest.

