## Supplemental Information for "Microbial Modulation of Host Plant Proteasome Activity Improves Heat Stress Tolerance"

**This PDF file includes:**

Figures S1 to S4
Tables S1 to S2
Dataset S1 (separate file)

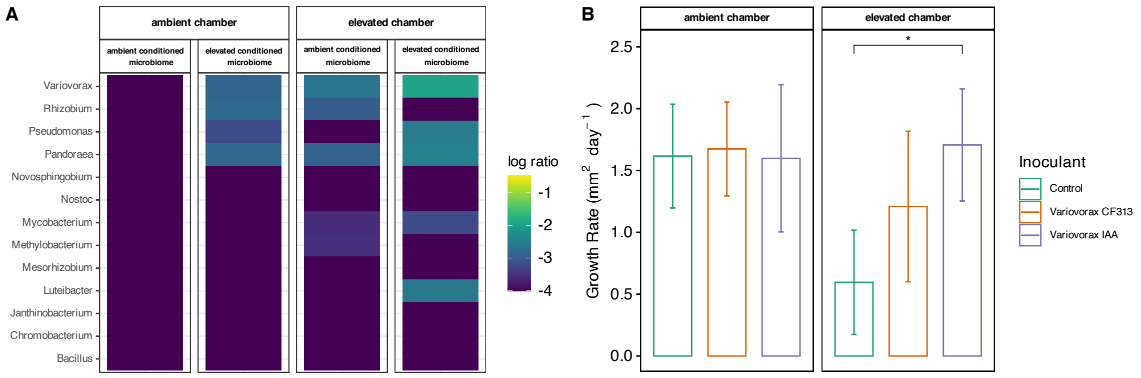


**Figure S1. *Variovorax* abundance and growth enhancement under ambient and elevated temperatures. (A)** Heatmap showing relative abundance of microbial taxa that match isolates in our culture collection, across experimental conditions. Columns are grouped by chamber temperature (ambient or elevated) and microbiome source (ambient- or elevated-conditioned field samples), with log-transformed abundance indicated by color intensity. **(B)** Growth rates of Sphagnum inoculated with *Variovorax* strains. Plants were inoculated with either *Variovorax* sp. CF313 (isolated from *Populus*), *Variovorax* IAA (isolated from SPRUCE), or a no-bacteria control, then grown under ambient or elevated temperatures. Points represent individual measurements, bars show means ± SD (n = 6 per treatment). Asterisks indicate significant differences between treatments ($P$<0.05, Wilcoxon rank-sum test with Bonferroni correction).

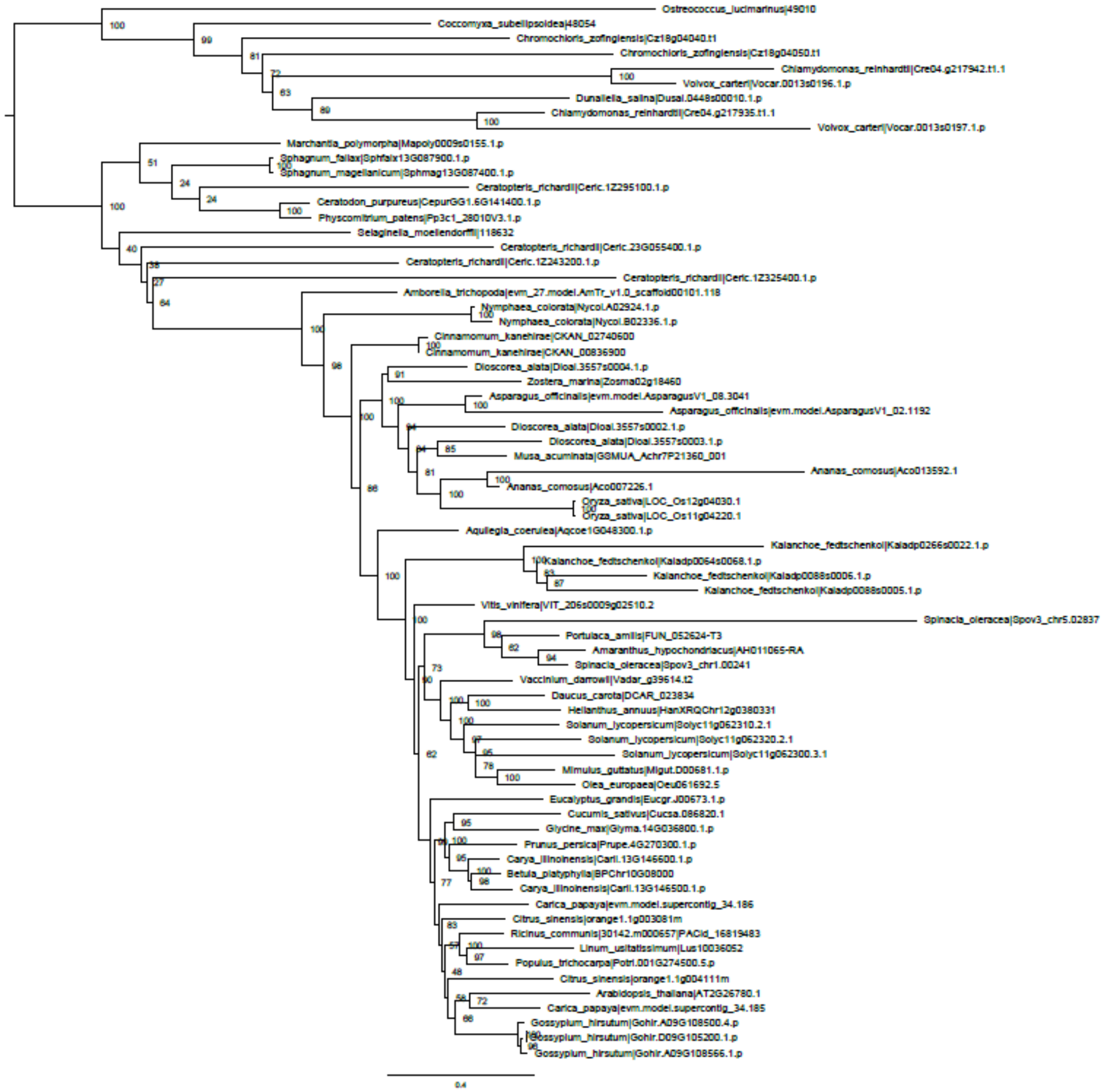


**Figure S2. Phylogenetic tree for *ECM29* across plant species, from red algae to angiosperms.**

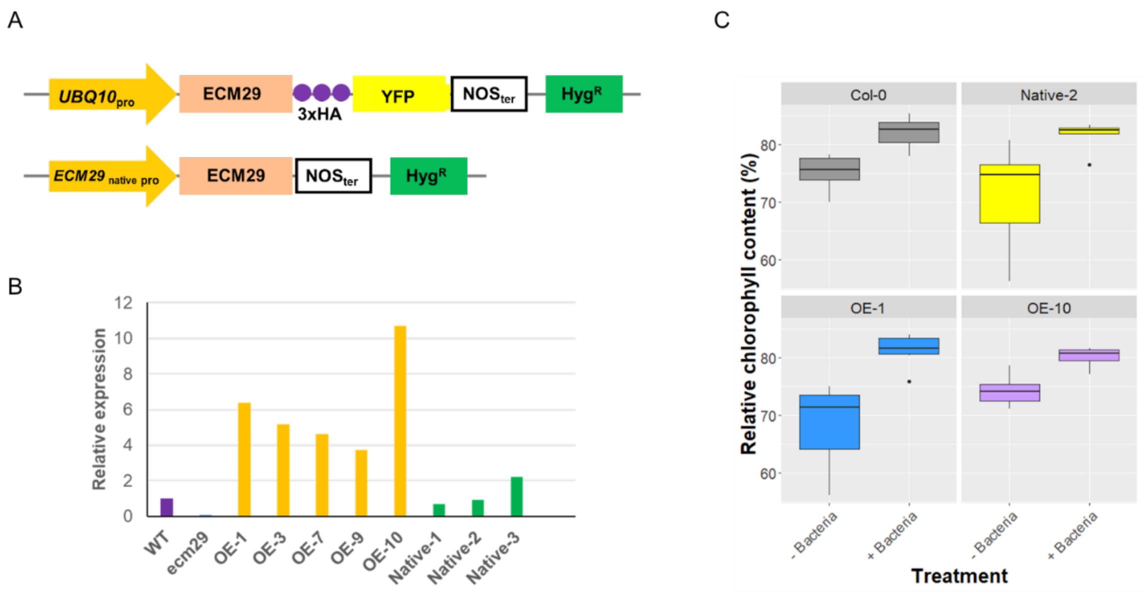


**Figure S3. Transgenic lines expressing *ecm29*. (A)** Schematic diagram of *ecm29* expressing cassettes driven by AtUBQ10 promoter or *ecm29* own promoter. **(B)** *ecm29* transcript expression profile in wilt-type, *ecm29* mutant, and transgenic plants. **(C)** Complementation of *ecm29* recovers mutant phenotype.

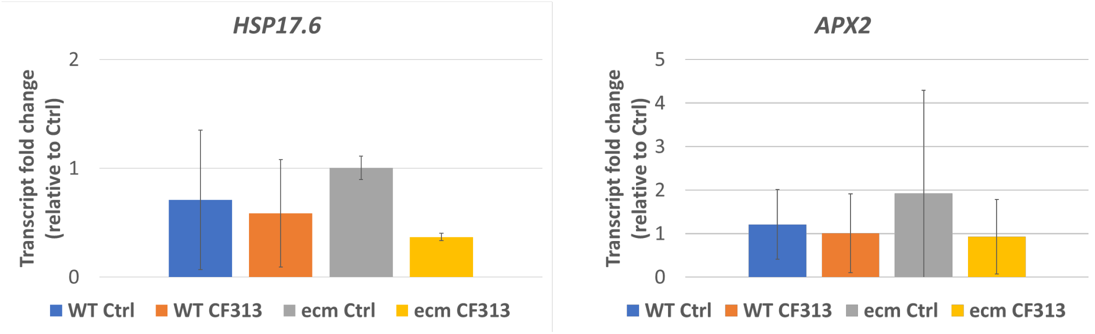


**Figure S4. Relative gene expression levels of heat-related genes in response to *Variovorax* sp. CF313.** *WT,* Col-0 wild-type; *ecm*, *ecm29* mutant; *Ctrl,* mock control without bacteria; *CF313,* inoculated with *Variovorax* sp. CF313.

**Table 1.** List of 26S proteasome subunits that interact with ECM29.

| | | Always | Inoculated |
|---|---|---|---|
| 19S RP Lid | | RPN5 | RPN6 |
| | | | RPN8 |
| | | | RPN12 |
| | Base | RPN1 | RPN10 |
| | | RPT2 | RPT1 |
| | | RPT3 | |
| | | RPT4 | |
| | | RPT5 | |
| | | RPT6 | |
| 20S CP | | | PAC |
| | | | PAD |

**Table S1.** List of candidate gene models for QTL analysis.

| Locus ID | Protein | Best-hit-*A.thaliana* | Function | Ref. |
|---|---|---|---|---|
| Sphfalx13G087500 | CN_hydrolase | | Hydrolysis of carbon-nitrogen bond | |
| Sphfalx13G087600 | RETINALDEHYDE BINDING PROTEIN-RELATED | AT1G01630 | Root hair development | (1) |
| Sphfalx13G087700 | PHOSPHOGLYCERATE MUTASE | AT2G17280 | Fungal elicitor response | (2) |
| Sphfalx13G087800 | GLYCOSYLTRANSFERASE 14 FAMILY MEMBER | AT5G39990 | Arabinogalactan biosynthesis | (3, 4) |
| Sphfalx13G087900 | PROTEASOME-ASSOCIATED PROTEIN ECM29 HOMOLOG | AT2G26780 | Proteasome quality control and assembly | (5, 6) |
| Sphfalx13G088000 | RHO FAMILY GTPASE | AT4G35020 | Pavement cell morphogenesis, cell signaling | (7–9) |
| Sphfalx13G088100 | ACTIN-RELATED PROTEIN 2/3 COMPLEX SUBUNIT 1 | AT2G30910 | Trichome morphogenesis | (10) |

**Table S2.** Conditioned microbiome laboratory incubation temperature and light cycle. Temperatures were determined from June average temperature for 6-hour blocks at the SPRUCE field site that microbiomes were isolated from.

| Growth Chamber Treatment | Time (hr) | Temperature (°C) | Light |
|---|---|---|---|
| Ambient | 0:00 | 13 | no |
| | 6:00 | 18 | yes |
| | 12:00 | 21 | yes |
| | 18:00 | 15 | no |
| Elevated | 0:00 | 22 | no |
| | 6:00 | 27 | yes |
| | 12:00 | 30 | yes |
| | 18:00 | 24 | no |
